# High feature overlap reveals the importance of anterior and medial temporal lobe structures for learning by means of fast mapping

**DOI:** 10.1101/2020.08.06.240697

**Authors:** Ann-Kathrin Zaiser, Regine Bader, Patric Meyer

**Affiliations:** Experimental Neuropsychology Unit, Department of Psychology, Saarland University, Saarbrücken, Germany; Department of Applied Psychology, SRH University Heidelberg, Germany; Network Aging Research (NAR), Heidelberg University, Heidelberg, Germany

**Author notes:** Corresponding author. Neurocognitive Psychology Unit, Department of Applied Psychology, SRH University Heidelberg, Maria-Probst-St. 3, 69123 Heidelberg, Germany. E-mail address (A.-K. Zaiser).

## Abstract

Contrary to traditional theories of declarative memory, it has recently been shown that novel, arbitrary associations can be rapidly and directly integrated into cortical memory networks by means of a learning procedure called fast mapping (FM), possibly bypassing time-consuming hippocampal-neocortical consolidation processes. In the typical FM paradigm, a picture of a previously unknown item is presented next to a picture of a previously known item and participants answer a question referring to an unfamiliar label. It is assumed that they thereby incidentally create associations between the unknown item and the label. However, contradictory findings have been reported and factors moderating rapid cortical integration through FM yet need to be identified. In the context of previous behavioral results showing rapid semantic integration through FM especially if the unknown and the known item shared many features, we propose that due to its computational mechanisms during the processing of complex and particularly highly similar objects, the perirhinal cortex might be especially qualified to support the rapid incorporation of these associations into cortical memory networks within the FM paradigm. We therefore expected that a high degree of feature overlap between the unknown and the known item would trigger strong engagement of the perirhinal cortex at encoding, which in turn might enhance rapid cortical integration of the novel picture-label associations. Within an fMRI experiment, we observed stronger activation for subsequent hits than misses during encoding in the perirhinal cortex and an associated anterior temporal network if the items shared many features than if they shared few features, indicating that the perirhinal cortex indeed contributes to the acquisition of novel associations by means of FM if feature overlap is high.

## 1. Introduction

Traditional theories of declarative memory assume that learning of novel, arbitrary associations depends on a time-consuming consolidation process, typically based on hippocampal-neocortical interplay (e.g., Frankland & Bontempi, 2005; McClelland, McNaughton, & O’Reilly, 1995). However, there is evidence that rapid and direct cortical integration of novel picture-label associations is possible by means of an encoding procedure called *fast mapping* (FM; e.g., Himmer, Müller, Gais, & Schönauer, 2017; Merhav, Karni, & Gilboa, 2014, 2015; Sharon, Moscovitch, & Gilboa, 2011; Zaiser, Meyer, & Bader, 2019b). Sharon and colleagues (2011) reported a clear benefit from encoding within the FM paradigm in patients with severe lesions predominantly to the hippocampus. Whereas these patients did not recognize novel picture-label associations above chance level after encoding through a standard explicit encoding (EE) condition that is typically expected to rely on hippocampal processing, their recognition performance was as good as that of healthy controls if the associations had been encoded by means of FM. In the typical FM paradigm, learning is incidental and thus, participants are not informed about later memory tests. They are presented with a picture of a previously unknown item (e.g., an exotic blue-footed bird) together with a picture of a previously known item that is already represented in semantic networks (e.g., a flamingo), and are asked to answer a question referring to an unfamiliar label (e.g., *Does the satellote have blue feet?*). Participants can answer this question by recognizing and rejecting the previously known item, thereby actively discovering the link between the picture of the unknown item and the unfamiliar label. It is assumed that this procedure enables the binding of the picture of the unknown item and the label to a new association that can be rapidly integrated into semantic memory networks (Sharon et al., 2011).

Despite evidence that FM can enable direct integration of arbitrary associations, other studies revealed contradictory findings (cf. Cooper, Greve, & Henson, 2019; Greve, Cooper, & Henson, 2014; Smith, Urgolites, Hopkins, & Squire, 2014; Warren & Duff, 2014; Warren, Tranel, & Duff, 2016). However, the experimental designs and procedures of some of these studies deviated from the original paradigm, such that learning was intentional (e.g., Warren & Duff, 2014) or the associations had been repeatedly recalled before the recognition memory test (e.g., Warren & Duff, 2014; Warren et al., 2016). Moreover, rapid cortical integration through FM was not always investigated in patients with lesions confined to the hippocampus but also in patients with extended lesions to extra-hippocampal structures or with complete left-temporal lobectomies (Warren et al., 2016). In addition, it is difficult to draw conclusions on rapid cortical integration from studies in which solely behavioral recognition tests have been used to assess retrieval from cortical networks in healthy young adults (e.g., Cooper et al., 2019) as these explicit tests alone do not allow for the dissociation between retrieval of hippocampal and cortical memory representations in such samples. A recent debate has underpinned the necessity to clarify more systematically if and under which conditions rapid cortical integration of novel associations through FM is possible (see Cooper, Greve, & Henson, 2018, and the respective commentaries), for example by identifying factors potentially moderating FM learning success (see Zaiser, Meyer, & Bader, 2019a). Here, we approached this issue from a neurocognitive perspective, asking which underlying mechanisms and corresponding brain structures are likely to contribute to successful rapid cortical integration through FM.

Apart from the benefit of encoding through FM for patients with lesions predominantly to the hippocampus, Sharon et al. (2011) showed that two additional patients who exhibited extended lesions to other temporal lobe structures, such as the perirhinal cortex (PrC) and the anterior temporal lobe (ATL), did not benefit from encoding through FM. There is a large body of evidence that the PrC as a key component of an anterior temporal system (see Ranganath & Ritchey, 2012) is involved in the processing and discrimination between complex objects, especially if they share many features (e.g., Bussey, Saksida, & Murray, 2005; Cowell, Bussey, & Saksida, 2010). For example, Barense, Gaffan, and Graham (2007; see also Barense et al., 2005) found that, in contrast to patients with lesions confined to the hippocampus, patients with lesions extending to the PrC could not discriminate between highly similar objects despite normal performance in the discrimination between less similar objects. As the discrimination between complex objects (i.e., the previously known and the unknown item) is one of the most central cognitive operations required in the FM encoding task, one could assume that the PrC is involved in encoding through FM and that this involvement should increase with enhanced similarity between the items. Interestingly, in the study by Sharon et al. (2011), the two pictures in the FM encoding screen were highly similar in order to make the task more demanding and thereby allow for deeper encoding (see also Sharon, 2010). Such a high feature overlap between the previously unknown and the known item might have triggered PrC-mediated processes during FM encoding, from which selectively patients with hippocampal but not with additional perirhinal lesions might have benefitted.

In addition to its perceptual role, the PrC is involved in semantic processing (e.g., Meyer et al., 2013; Meyer, Mecklinger, & Friederici, 2010; Meyer et al., 2005; Wang, Lazzara, Ranganath, Knight, & Yonelinas, 2010; Wang et al., 2014) and familiarity-based item recognition memory (e.g., Bowles et al., 2007; Bowles et al., 2010; see Brown & Aggleton, 2001, for a review). A representational-hierarchical view of the medial temporal lobe suggests that the PrC generally processes complex conjunctions of elemental features as single units, irrespective of the domain (i.e., on a perceptual as well as a semantic and mnemonic level; Cowell, Barense, & Sadil, 2019; Cowell, Bussey, & Saksida, 2006; O’Neil, Barkley, & Köhler, 2013). This cross-domain role of the PrC suggests that discriminative and mnemonic factors might interact during encoding. In line with this assumption, Chen, Zhou, and Yang (2019) found that the increased activation of the PrC during the discrimination between two highly similar items at encoding, compared to the discrimination between items from different categories, was predictive of later item recognition memory. Supportive evidence comes from an eye-tracking study by Zhou, Chen, and Yang (2018), who found that more saccades between similar items were also predictive of subsequent item memory. In addition to the processing of single items, familiarity-based memory for newly built associations between items was found to be accompanied with enhanced PrC contribution to learning if the associations are encoded as integrated units (Haskins, Yonelinas, Quamme, & Ranganath, 2008). Here, we set out to ask if the PrC could similarly support memory processes in learning of associations through FM, such that stronger PrC recruitment at encoding – triggered through increased demands on the discrimination between the known and unknown item – would support the binding of the novel association to a unit, thereby potentially facilitating their rapid incorporation into cortical networks.

Hence, here we explicitly manipulated the demands on the discrimination between the previously known and the unknown item (as in Zaiser et al., 2019b) in an fMRI experiment, contrasting an FM encoding condition in which the previously unknown and the known item shared many features (*fast mapping, high feature overlap*; FMHO) to an FM encoding condition in which they shared few features (*fast mapping, low feature overlap*; FMLO). We expected that PrC engagement at encoding should be greater if the demands on perirhinal processing (i.e., the discrimination between complex objects) are higher (as in the FMHO condition) than if they are lower (as in the FMLO condition). We assume that this should also be reflected in differential PrC contribution to learning. In particular, especially in the FMHO condition, stronger PrC involvement was expected at encoding of items that were subsequently remembered in a forced-choice recognition test than at encoding of items that were subsequently forgotten (i.e., larger subsequent memory effects in the FMHO condition compared to the FMLO condition). If PrC engagement drives rapid cortical integration by means of FM and this can be enhanced by increasing feature overlap, this could pave the way to hippocampus-independent learning of novel, arbitrary associations and might explain why hippocampal consolidation could be bypassed in an FM study using high feature overlap pairs (Sharon et al., 2011).

## 2. Method

### 2.1 Participants

Data were collected until 48 complete datasets of healthy participants showing above-chance recognition performance were obtained. Participants were pseudo-randomly assigned to an FMHO and an FMLO group until both groups contained 24 participants (FMHO: *M*_age_ = 24.1 years, age range: 19-30; FMLO: *M*_age_ = 22.1 years, age range: 18-26) and gender distribution was the same in both groups (14 female each). All participants were right-handed in accordance with the Edinburgh Handedness Inventory (Oldfield, 1971) and native German speakers. Of the total sample of *N* = 97 participants, 13 were excluded due to arachnoid and pineal cysts, one participant due to a panic attack in the scanner, and one participant as he had already taken part in another experiment using the same materials. Further three participants were excluded as not enough trials (< 10) remained after exclusion of trials based on a post-experimental rating of prior knowledge (see Design and Procedure section). Of the remaining 79 participants, further 31 participants were excluded from the analyses as they did not show above-chance recognition accuracy (*p* > .05, binomial test; *n*_FMHO_ = 23; *n*_FMLO_ = 8). Participants gave written informed consent prior to the experiment and were compensated for their participation with 8€ per hour. The experiment was approved by the ethics committee of the Faculty of Human and Business Sciences at Saarland University in accordance with the declaration of Helsinki.

### 2.2 Materials

All pictures were drawn from the internet and arranged in pairs of one putatively known and one putatively unknown item each. For reasons of counterbalancing between encoding conditions, each unknown item was assigned a highly similar known item (for usage in the FMHO condition) and a less similar known item (for usage in the FMLO condition; see Figure 1). Analogously, each known item could appear together with one highly similar and one less similar unknown item. Single items belonged to one of seven categories (mammals, birds, insects, fish, fruit, vegetables, plants). In a previously conducted rating study, a different sample of 46 participants had rated these items for familiarity (5-point Likert scale; 1 = *not at all familiar*, 5 = *very familiar*) and previous knowledge (*known* vs. *unknown*). Item pairs, consisting of one putatively unknown and one putatively known item, had been rated for feature overlap, which was defined as the number of features the two pictures have in common (e.g., the presence and nature of fur, a tail, legs, the similarity of colors, etc.) and was rated on a 5-point Likert scale (1 = *not at all similar*, 5 = *very similar*).

**Figure 1.**
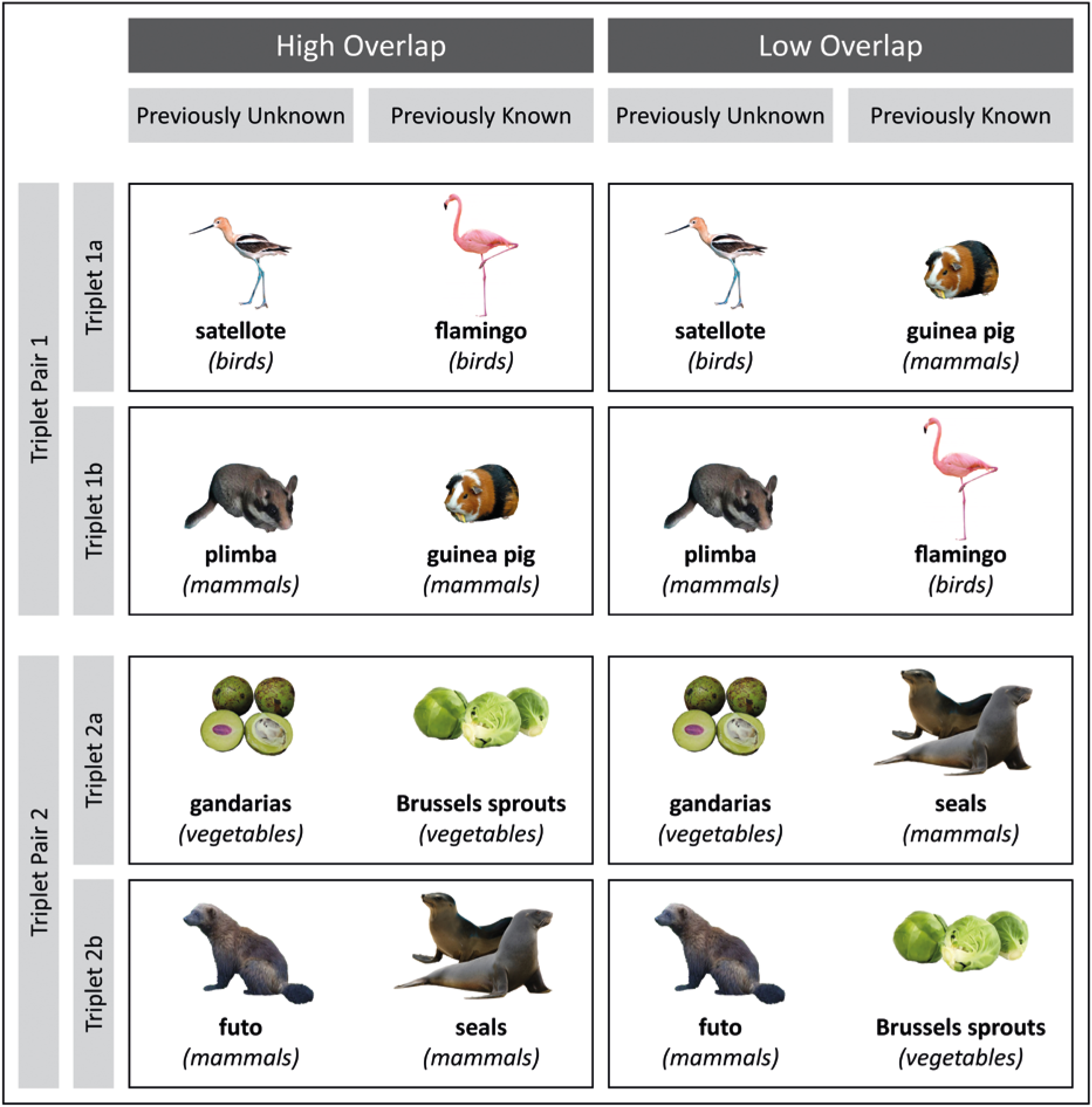
Example stimulus material. Each line depicts a picture triplet, consisting of one previously unknown item and two previously known items. Triplets were arranged in triplet pairs (e.g., Triplet Pair 1: Triplet 1a and 1b), within which overlap of the unknown and known items was counterbalanced. One of a triplet’s previously known items was for the high-overlap encoding condition (e.g., Triplet 1a: flamingo) and one for the low-overlap encoding condition (e.g., Triplet 1a: guinea pig). Overlap condition of the previously known item was interchanged in the other triplet of a triplet pair (e.g., flamingo as low-overlap known item and guinea pig as high-overlap known item in Triplet 1b). High-overlap item pairs were always from the same lower-level category (e.g., Triplet 1a: both birds). Low-overlap item pairs could consist of items from the same higher-level category but different lower-level categories (Triplets 1a and 1b: both animals, with birds and mammals as lower-level categories) or from different higher-level categories (Triplets 2a and 2b: plants and animals, with vegetables and mammals as lower-level categories).

Forty-eight item triplets, consisting of one unknown item, its highly similar known item, and its less similar known item (see Figure 1), were drawn from the stimulus material of the rating study. The counterpart of a triplet, that is, the triplet in which the putatively known items appeared in the respective other overlap condition, was also included. Hence, each unknown and known item appeared within a high-overlap pair in the FMHO group and within a low-overlap pair in the FMLO group. Of the triplets included in the present study, the previously unknown item had been classified as unknown by most participants in the rating study (on average, by 90 %, *SD* = 12 %) and had been rated with the lowest familiarity (*M* = 2.08, *SD* = 0.43), and the previously known items had been rated as known by most participants (on average, by 85 %, *SD* = 12 %) and with the highest familiarity (*M* = 4.42, *SD* = 0.40). Moreover, only triplets with the highest difference between the overlap rating of the high-overlap and the low-overlap item pair were included (*M*_FMHO_ = 3.59, *SD*_FMHO_ = 0.51; *M*_FMLO_ = 1.42, *SD*_FMLO_ = 0.37; *M*_diff_ = 2.17, *SD*_diff_ = 0.62). In the final item set, significantly more participants of the rating study had rated the previously known item as known than the previously unknown items, familiarity for the previously unknown items was significantly lower than for the previously known items, and overlap of the high feature overlap pairs was higher than overlap of the low feature overlap pairs (all *p*s < .001). In addition, the lowest overlap rating of the high-overlap pairs was still higher than the highest overlap rating of the low-overlap pairs. Further 12 trials were added as filler trials, in which the question referred to the previously known item, which was supposed to prevent participants from developing strategies such as always referring to the unknown item without paying attention to the known item. Filler trials matched the participants’ encoding condition with regard to feature overlap and were excluded from all analyses.

Half of the questions at encoding required a positive response, half a negative response, and questions were identical for both overlap conditions (e.g., the question *Does the satellote have blue feet?* was asked no matter if the satellote was paired with the highly similar flamingo or the less similar guinea pig). The items’ actual names were substituted with their botanical or zoological name (sometimes slightly modified) or with a pseudo-word if these labels might have triggered expectations about an item’s category or features (e.g., if the name contained information on the item, such that *giraffe gazelle* would indicate a hoofed animal and was thus given its zoological name *gerenuk*). Word length of all labels was between 4 and 10 letters (*M* = 6.88, *SD* = 1.84).

### 2.3 Design and procedure

Stimulus presentation and timing were controlled using the experimental software PsychoPy (Peirce, 2009; http://www.psychopy.org/). All stimuli throughout the experiment were presented against a white background, projected onto a screen behind the magnet which was visible through a mirror attached to the head coil. All tasks except for the post-experimental stimulus rating were conducted in the scanner; the encoding and recognition phase were recorded. Responses were collected via two 2-button response grips (one in each hand), with which participants could respond by pressing one of two buttons on either side (left and right thumb and index finger).

#### Encoding

In order to ensure incidental learning, participants were told that visual perception would be investigated. All participants encoded the same picture-label associations by means of FM and feature overlap was manipulated between subjects. They first completed six practice trials (including two filler trials with the question referring to the previously known item), matching their individual overlap condition. In the actual encoding phase, 60 experimental trials (including 12 filler trials) were presented in random order with the constraint that one of the filler trials was presented at the beginning and one at the end of the encoding phase in order to prevent primacy and recency effects. Each trial started with a fixation cross that was horizontally centered and slightly below the center of the screen, at the same height as the question would subsequently appear. The duration of this inter-stimulus interval was jittered between 1000 and 8000 ms in equally distributed steps of 500 ms. After the fixation cross had disappeared, the question was displayed separately for the first 2000 ms in Arial 27 point font and together with the pictures for further 3500 ms (see Figure 2). The label within the question was presented in the horizontal center of the screen in bold font. Participants were instructed to read the question thoroughly and, as soon as the pictures would appear, to identify the item to which the question refers and how it is thus to be answered. After both the pictures and the question had disappeared, the words *yes* and *no* were displayed on the left and right side of the screen in orange and blue color (position and color counterbalanced between subjects). Responses could be made by pressing the keys at the left or right index finger on the response grips. After 3000 ms, participants received written verbal feedback and moved on to the next trial. If no answer had been given within this time, they were encouraged to respond faster. In contrast to most previous FM studies, we decided against the repetition of encoding trials as repeating the associations would have prevented from capturing the effects of one-shot learning.

**Figure 2.**
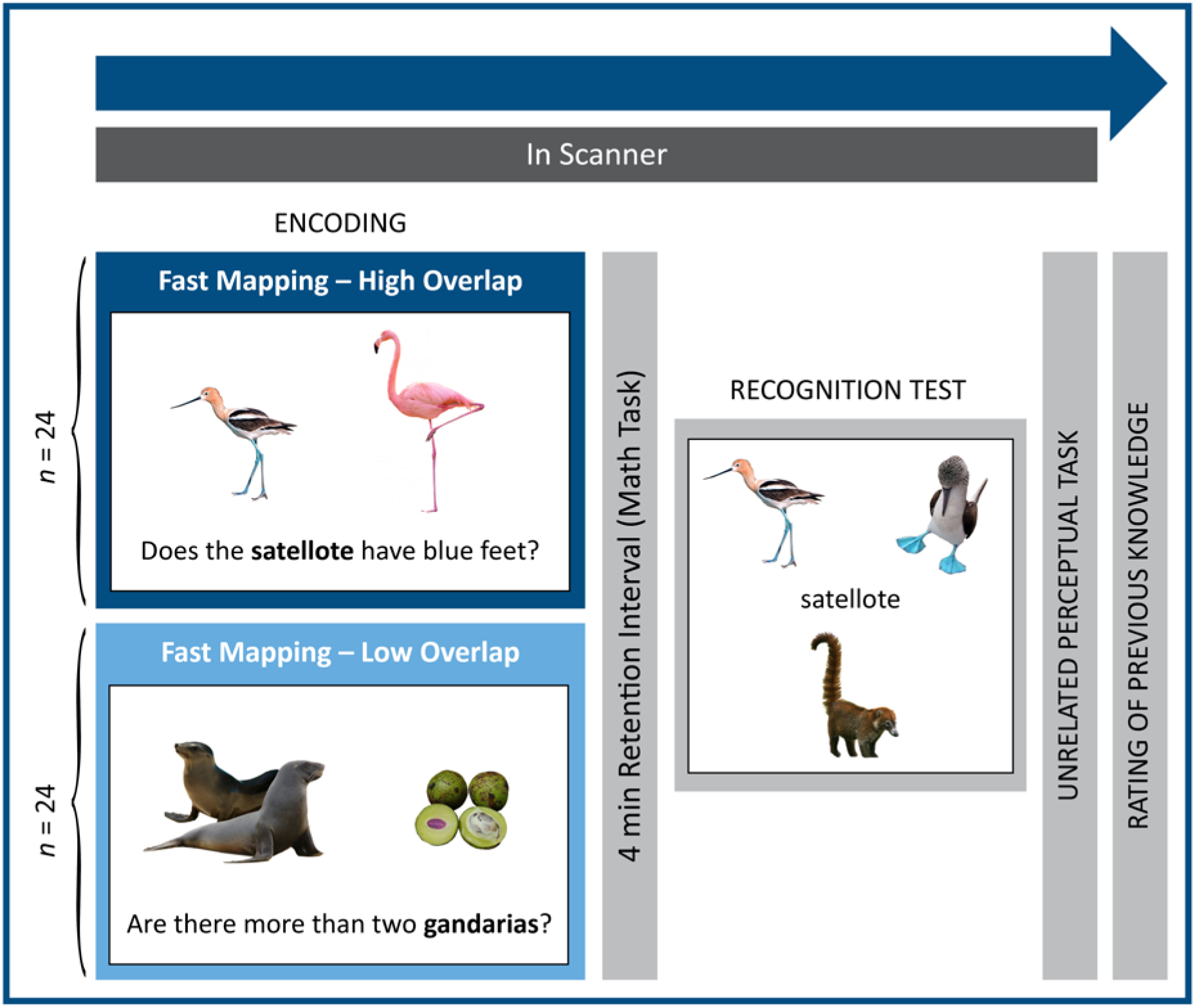
Experimental design and procedure. Encoding condition was manipulated between subjects. The plain question was presented for 2000 ms and then together with the pictures for further 3500 ms. After the pictures and the question had disappeared, response options (*yes*/*no*) were displayed and feedback was given after a response had been made. At recognition, the target and two foil pictures within one display always belonged to the same higher-level category (i.e., all animals or all plants) and had all been presented in the encoding phase.

#### Recognition

After a 4-min filler task, in which participants had solved simple mathematical equations, a three-alternative forced-choice recognition test was administered in which participants were tested for all 48 picture-label associations. A fixation cross was displayed in the center of the screen for a jittered interval between 1000 and 8000 ms in equally distributed steps of 500 ms (with 1000 ms, 4500 ms, and 8000 ms appearing four times), before it was replaced by the recognition test label (see Figure 2). The target picture and the two foil pictures were arranged around the label, with their positions on the screen randomly assigned (top-left, top-right, bottom-center). Participants were instructed to indicate which of the three pictures belonged to the test label by pressing the respective button on the response grips (left thumb, right thumb, right index finger). All three pictures had appeared in the encoding phase and were always from the same higher-level category (i.e., both animals or both plants) in order to control for item familiarity. Responses could not be given before 3000 ms of stimulus presentation, indicated by a verbal prompt at the bottom of the screen, in order to ensure sufficient exposure time to all pictures. The next trial started after 6000 ms of overall stimulus presentation. No feedback was provided. Prior to the actual recognition test, participants had completed a practice phase of four trials in which the four novel associations of the encoding practice phase were tested. After completion of the recognition task, an unrelated perceptual task was administered.

#### Rating of previous knowledge

Outside the scanner, participants’ individual prior knowledge of all items was assessed. Participants were seated in front of a 17-inch laptop at a viewing distance of approximately 50 cm, where they were informed that the main aim of the experiment was to investigate memory and it was necessary to assess which items they had already known prior to their participation. They were also informed that the stimuli were renamed and they were asked to indicate prior knowledge irrespective of an item’s label in the experiment. The participants then were sequentially presented with all pictures in random order and were instructed to rate how well they had known each item prior to the experiment on a 5-point Likert scale (1 = *had not known the item at all before the experiment*; 5 = *had known the item very well before the experiment*). After ratings of >= 4, they were asked to type in the item’s name at the lowest category level possible (e.g., *hawk* instead of *bird*).

### 2.4 Data acquisition and processing

A 3T Siemens Magnetom Skyra scanner with a 20-channel head coil was used for structural and functional data acquisition. Structural data were acquired prior to the experiment, using a T1-weighted three-dimensional magnetization-prepared rapid gradient-echo sequence (TR = 1900 ms; TE = 2.13 ms; sagittal orientation; flip angle = 9°; voxel size = 0.9 mm isotropic, distance factor = 50%; 192 slices; FoV = 240 mm). For the functional scans, a T2*-weighted two-dimensional gradient-echo planar imaging sequence (TR = 2000 ms; TE = 30 ms; flip angle = 90°; voxel size = 3 mm isotropic; distance factor = 25%; matrix = 64 x 64; FoV = 192 mm; right-left phase encoding direction) was used for both runs (one at encoding, one at recognition). Thirty-six transversal slices were acquired (interleaved, ascending), oriented in parallel with the anterior-posterior commissure plane and then tilted by −30° (anterior upward) in order to reduce susceptibility artifacts in anterior and medial temporal lobe structures (see e.g., Weiskopf, Hutton, Josephs, & Deichmann, 2006). Before scanning, it was made sure that the FoV covers all regions of interest. In order to allow for signal equilibrium, the first four volumes of each functional run were discarded.

Imaging data were processed using SPM 12 (https://www.fil.ion.ucl.ac.uk/spm/software/spm12/). The 398 volumes of the encoding phase and 260 volumes of the recognition test phase were corrected for slice acquisition time using the first slice of each volume as reference image. They were motion-corrected by realignment of all images of a run to its first image and then co-registered to each participant’s anatomical T1 image. After segmentation into gray and white matter, cerebrospinal fluid, bone, soft tissue, and air, they were spatially normalized to the Montréal Neurological Institute (MNI) standard T1 template with interpolation to 2-mm isotropic voxels and then smoothed using a Gaussian 7-mm full-width half-maximum kernel. Images were visually inspected for artifacts and adequacy of motion correction and transformation into standard space.

### 2.5 Analyses

Analyses were conducted using R (R Core Team, 2016) and SPM 12 (https://www.fil.ion.ucl.ac.uk/spm/software/spm12/) for imaging data. For all analyses, trials were only included if the individual rating of prior knowledge was congruent with what was expected at stimulus creation, that is, if the putatively unknown item was classified as unknown by a participant (i.e., a rating of prior knowledge of <= 3) and if the putatively known item was classified as known (i.e., a rating of >= 4). Neither the number of remaining subsequently remembered trials (*M* = 21.75; range: 17-29) differed between overlap groups, *t*(46) = −1.99, *p* =. 278, nor the number of subsequently forgotten trials (*M* = 17.13; range: 10-23), *t* < 1. Participants with less than 10 remaining knowledge-congruent trials for at least one subsequent-memory condition (subsequently remembered, subsequently forgotten) were removed from the sample and replaced by new participants (*n* = 3).

#### 2.5.1 Behavioral analyses

Encoding and recognition accuracy represents the percentage of correct responses. All *t* tests comparing performance between groups were two-tailed and significance level of all tests was set to α =. 05.

#### 2.5.2 fMRI analyses

Individual time series were modeled with separate regressors for subsequently remembered and subsequently forgotten trials in the encoding phase and for correct and incorrect trials in the recognition test phase. For each run, six motion parameters were added as regressors of no interest and a high-pass filter with a 128-seconds cutoff was applied. The regressors were created by convolving the stimulus function related to event onset (i.e., time of picture onset for both the encoding and the recognition run) with a canonical hemodynamic response function. One contrast image was computed for each subject and phase (encoding: subsequently remembered > subsequently forgotten; recognition: correct > incorrect). The contrast of subsequently remembered > subsequently forgotten trials at encoding will be referred to as *subsequent memory effects*. In order to investigate differential subsequent memory effects between the FMHO and FMLO group, a second-level difference of FMHO subsequent memory effects and FMLO subsequent memory effects will be referred to as *interaction contrast* in the following. The interaction contrast at recognition is the group difference between the contrasts between correct and incorrect trials. An explicit mask was applied covering the whole brain (constructed from the WFU Pickatlas toolbox 3.0.5; Maldjian, Laurienti, Kraft, & Burdette, 2003). Generally, the *p*-value threshold was set to *p* =. 001, uncorrected, and a minimum cluster size of 10 contiguous voxels was used for the analyses. The *p*-value threshold for analyses within the PrC, the hippocampus, and anterior temporal structures was set to *p* =. 005, uncorrected, at a minimum cluster size of five contiguous voxels, due to the lower signal-to-noise ratio as a consequence of susceptibility artefacts in the medial temporal lobe and adjacent structures (see e.g., Davachi and Wagner, 2002; Dobbins, Rice, Wagner, & Schacter, 2003; Ojemann et al., 1997; O’Kane, Insler, & Wagner, 2005; Schacter & Wagner, 1999; Staresina & Davachi, 2006; Strange, Otten, Josephs, Rugg, & Dolan, 2002). We defined the PrC as Brodmann area (BA) 36 and the ATL as BA 38 as well as 20 and 21 for clusters with peaks anterior to the most posterior part of BA 38 (*y* = 0), with BAs according to the WFU Pickatlas 3.0.5 (Maldjian et al., 2003).

## 3. Results

### 3.1 Behavioral results

On average, 92.90 % (*SD* = 5.42 %) of the questions in the encoding phase were answered correctly and the proportion of correct encoding trials did not differ between subsequently remembered and forgotten trials, *t*(47) = −1.38, *p* =. 174, neither in the FMHO condition, *t* < 1, nor in the FMLO condition, *t*(23) = −1.20, *p* =. 241, all two-tailed. In addition, the difference of the correctly answered encoding questions for subsequently remembered versus forgotten items was not different between the FMHO and FMLO group, *t* < 1. At recognition, participants successfully recognized *M* = 56 % (*SD* = 6 %) of the picture-label associations and recognition accuracy was not different between the FMHO group (*M* = 55 %, *SD* = 5 %) and the FMLO group (*M* = 57 %, *SD* = 6 %), *t*(46) = −1.40, *p* =. 169, two-tailed.

### 3.2 Imaging results

#### Encoding

In order to check if PrC activation during perception of highly similar versus dissimilar pictures was actually different, we first investigated if the PrC was generally recruited more strongly at encoding in the FMHO condition than in the FMLO condition, irrespective of subsequent memory success. This was the case in the left PrC, *t* = 3.81 (peak: *x* = −22, *y* = −10, *z* = −28; cluster size = 32 voxels), and in the right PrC, *t* = 3.23 (peak: *x* = 24, *y* = −18, *z* = −24; cluster size = 10 voxels).

In order to test our main hypothesis that PrC contribution to FM learning should be greater in the FMHO condition than in the FMLO condition, we compared subsequent memory effects (subsequently remembered trials > subsequently forgotten trials) for the encoding conditions (subsequent memory effect FMHO > subsequent memory effect FMLO). As expected, greater subsequent memory effects in the FMHO condition than in the FMLO condition were found in the right PrC, *t* = 4.51 (peak: *x* = 28, *y* = −12, *z* = −26; see Table 1 and Figure 3). Separate analyses for this cluster within each group showed that this interaction was driven by a positive subsequent memory effect for the FMHO condition, *t*(23) = 2.71, *p* =. 006, *d* = 0.55, and a negative subsequent memory effect for the FMLO condition, *t*(23) = −2.49, *p* =. 021, *d* = −0.51, two-tailed (see Figure 3). Moreover, engagement of the PrC in encoding of remembered items was greater in the FMHO condition than in the FMLO condition, *t*(46) = 2.83, *p* =. 003, *d* = 0.82, whereas no differences in PrC involvement between encoding conditions were observed for subsequently forgotten items, *t* < 1 (see Figure 3).

**Table 1.**
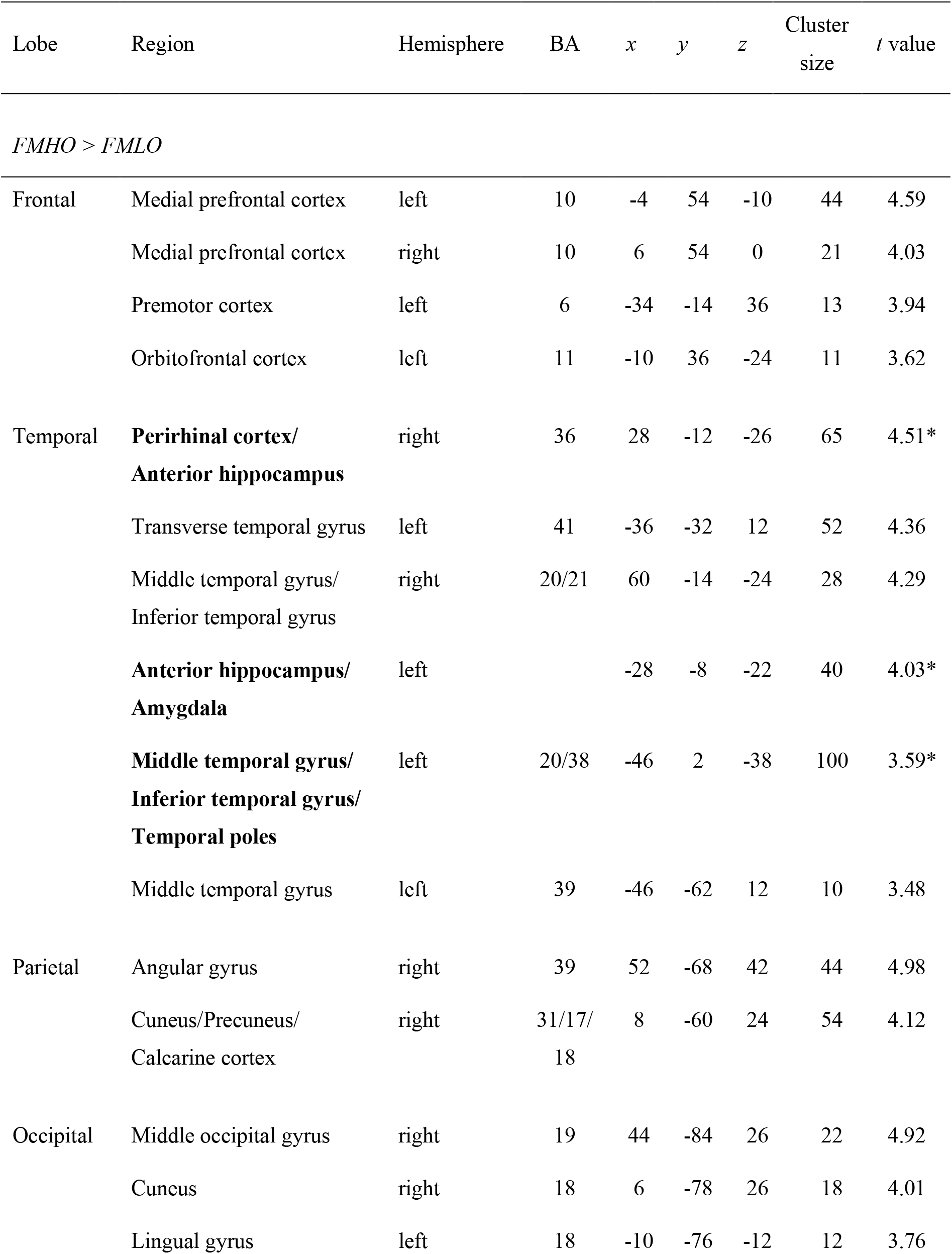
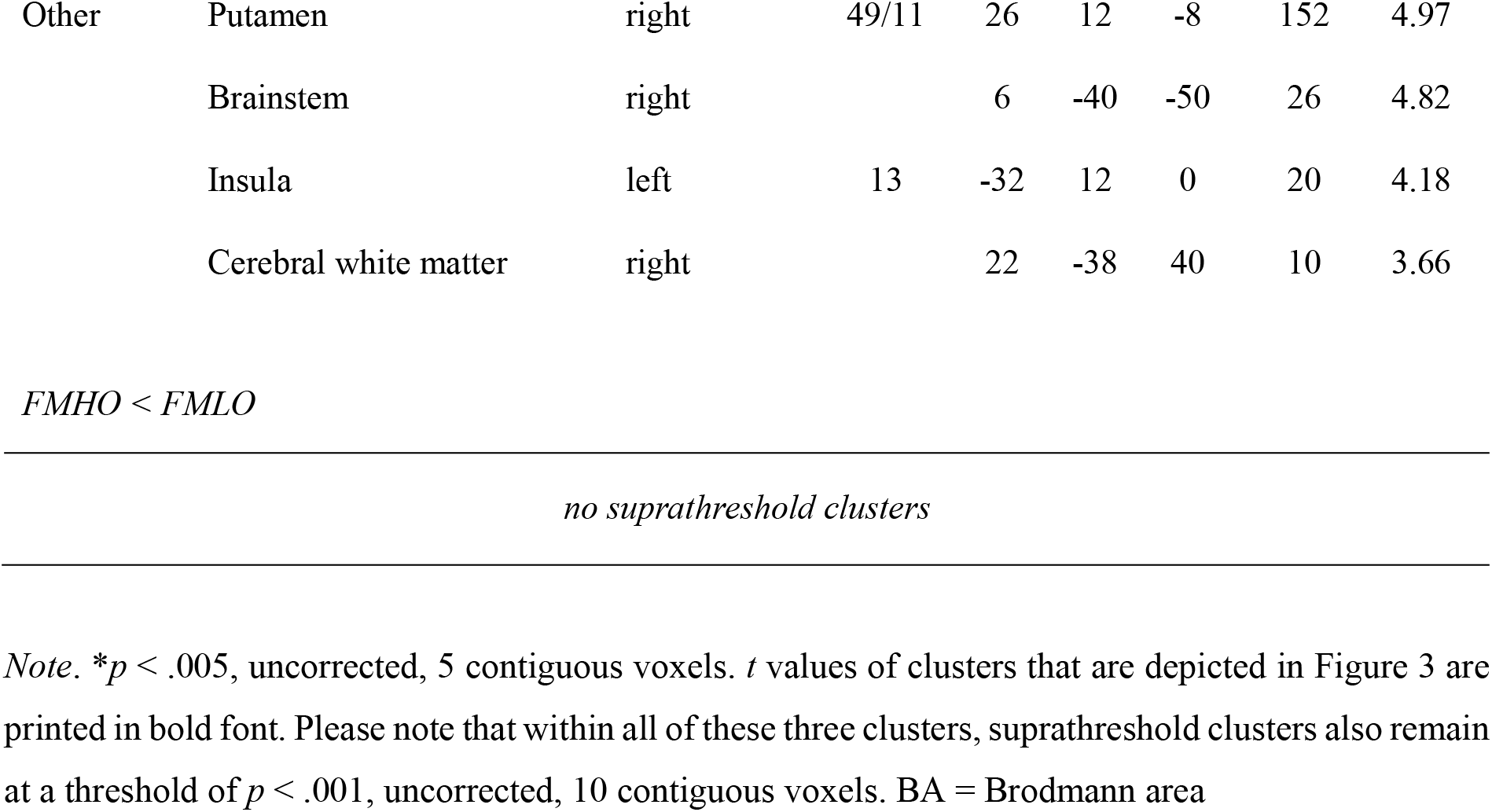
*Local Maxima of Clusters Showing Differential Subsequent Memory Effects Between Encoding Conditions, at p <. 001, Uncorrected*

**Figure 3.**
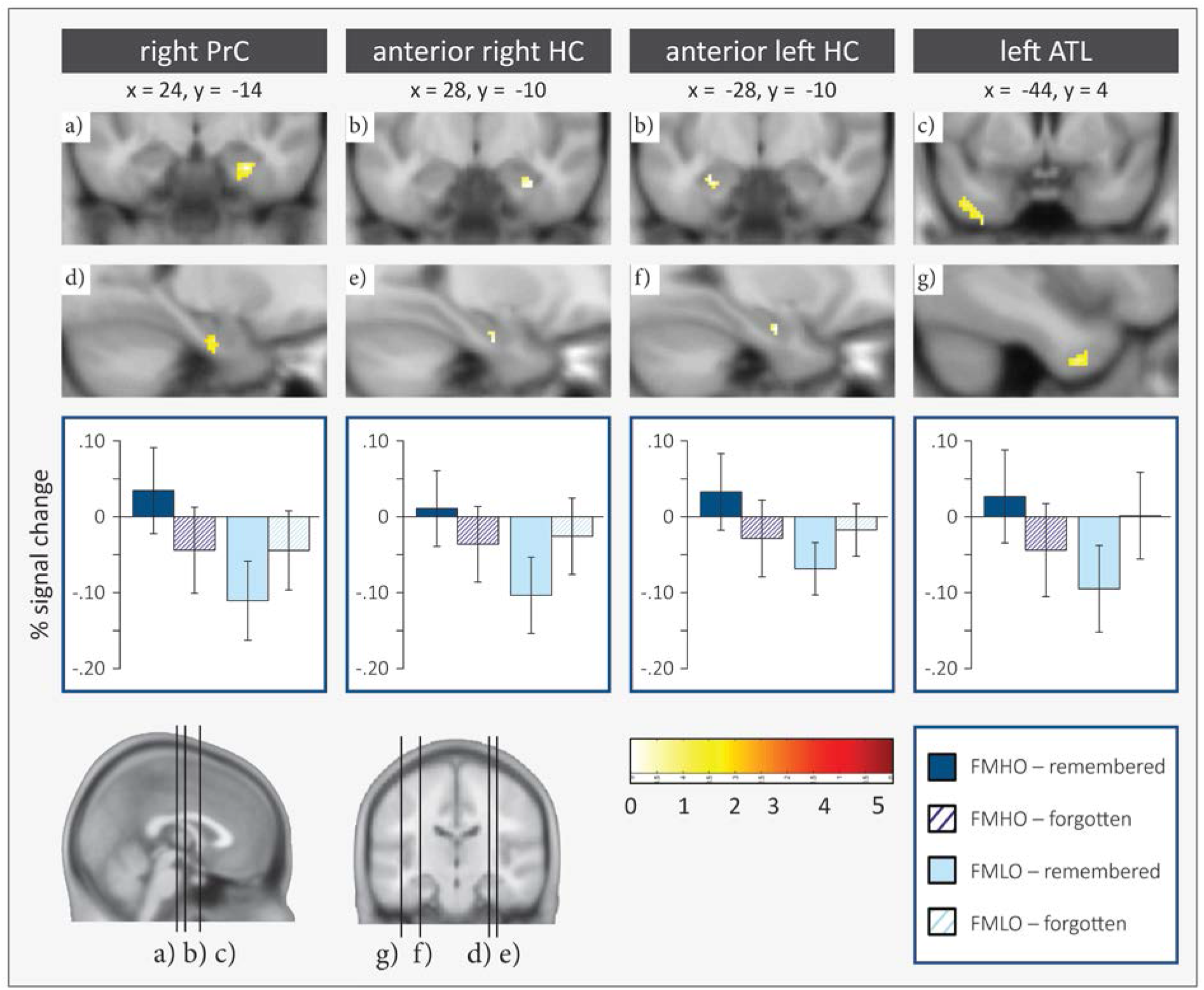
Selected clusters in which subsequent memory effects were observed to be greater for the FMHO condition compared to the FMLO condition. Error bars represent the two-tailed within-subjects confidence intervals of the difference between percent signal change at encoding of subsequently remembered compared to subsequently forgotten trials. Perirhinal cortex, hippocampus, and anterior temporal lobe clusters were determined using masks created with the WFU Pickatlas toolbox 3.0.5 (Maldijan et al., 2003; dilated by two voxels in three dimensions). PrC = perirhinal cortex, HC = hippocampus, ATL = anterior temporal lobe; FMHO = fast mapping, high overlap, FMLO = fast mapping, low overlap.

In addition to the contribution of the PrC to learning, the analyses revealed further clusters showing the interaction effect in, amongst others, the ATL and the anterior hippocampus bilaterally, the medial prefrontal cortex (mPFC), and the left orbitofrontal cortex (see Table 1). Notably, the patterns of signal change underlying the interaction effects in the regions named above are remarkably similar, that is, positive subsequent memory effects in the FMHO condition and negative subsequent memory effects in the FMLO condition (see Figure 3).

#### Recognition

Further analyses comparing involvement for correct vs. incorrect recognition trials at retrieval revealed that a cluster in the right PrC and right anterior hippocampus, *t* = 3.57 (peak: *x* = 22, *y* = −8, *z* = −28), cluster size = 24, seems to contribute to retrieval success (i.e., correct > incorrect) more in the FMHO condition than in the FMLO condition. The same interaction was identified in the ATL and in particular the right temporal pole, *t* = 4.21 (peak: *x* = 36, *y* = 6, *z* = −44), cluster size = 45, and the left temporal pole, *t* = 3.06 (peak: *x* = −36, *y* = 4, *z* = −42), cluster size = 24. The reverse interaction contrast, indicating larger effects for the FMLO compared to the FMHO group, was found in the left entorhinal cortex, *t* = 2.98 (peak: *x* = −18, *y* = −2, *z* = −34), cluster size = 8.

## 4. Discussion

There has been an extensive debate on the phenomenon of FM, questioning if FM enables rapid, direct cortical integration of novel associations, potentially bypassing the slow hippocampus-dependent consolidation processes that would typically be expected in memory for associations (e.g., Frankland & Bontempi, 2005; McClelland et al., 1995). We used a neurocognitive approach in order to identify factors that could potentially moderate learning within the FM paradigm and its underlying neurofunctional processes. One critical cognitive operation within the FM paradigm is the discrimination between the unknown and the known item. The PrC is especially qualified to be engaged in the discrimination between complex and highly similar objects (e.g., Mundy et al., 2012; Barense et al., 2005) as well as in memory for single items (see Brown & Aggleton, for a review) or unitized representations (Haskins et al., 2008). Consequently, we argued that high demands on the discrimination between the unknown and the known item in the FM paradigm might be associated with stronger PrC engagement, which might support the binding of the label to the unknown item. In a next step, this could facilitate the rapid and direct integration of this novel unit into cortical memory networks. We manipulated feature overlap between the unknown and the known item with the idea that the demands on perirhinal processing are especially high in the FMHO condition, which should recruit the PrC more strongly. We expected that this should lead to a stronger contribution of the PrC to learning in the FMHO condition compared to the FMLO condition. This was confirmed by the present experiment, revealing subsequent memory effects within the right PrC in the FMHO condition and greater PrC subsequent memory effects for the FMHO condition than for the FMLO condition.

There is complementary behavioral evidence that rapid semantic integration (as measured by means of semantic priming effects) through FM benefits from a high similarity between the objects that need to be discriminated at encoding (Zaiser et al., 2019b). However, despite this evidence for feature overlap as a moderating factor, it has not yet been investigated which underlying neurocognitive mechanisms and neurofunctional correlates are associated with learning by means of FM. Many previous FM studies point to the ATL as key candidate for rapid semantic integration through FM (e.g., Atir-Sharon, Gilboa, Hazan, Koilis, & Manevitz, 2015; Sharon et al., 2011; Merhav et al., 2015). This is reasonable insofar as the ATL has repeatedly been identified as a semantic hub, receiving input information from multiple modality-specific sensory areas which are then integrated into a coherent concept in the ATL (see e.g., see Lambon Ralph, Jefferies, Patterson, & Rogers, 2017; and Patterson, Nestor, & Rogers, 2007, for reviews). Ranganath and Ritchey (2012) suggested that the ATL is part of an anterior temporal system, one of two systems in their model for memory-guided behavior. Apart from anterior parts of the ATL, this anterior temporal system encompasses the lateral orbitofrontal cortex, the amygdala, anterior regions of the hippocampus, and, as a key component, the PrC. Atir-Sharon et al. (2015) observed that the ATL specifically contributes to learning by means of FM but not EE, and Merhav et al. (2015) reported the engagement of the ATL and ATL-related networks at retrieval of associations shortly after they had been acquired through FM (but not EE), potentially affording a direct route to cortical integration of the associations. Furthermore, the pattern of residual ATL volumes in the patients reported by Sharon et al. (2011) clearly distinguishes between the four amnesic patients who benefitted from FM and two other patients who showed no learning benefit. In addition, there is complementary fMRI evidence for ATL contribution to learning by means of FM (Atir-Sharon et al., 2015) and the involvement of ATL-related networks in retrieval of associations acquired through FM (Merhav et al., 2015). In contrast to the ATL, the role of the PrC in rapid cortical integration through FM has not been investigated explicitly, although in the Sharon study, PrC volume seems to correlate with recognition accuracy in the FM condition. Factors leading to differential PrC engagement during FM learning (e.g., feature overlap) have not been manipulated or controlled in previous studies. Moreover, as the PrC is prone to susceptibility artefacts in fMRI (see e.g., Weiskopf et al., 2006), the identification of PrC activation in some studies might have become even more difficult.

The present fMRI experiment showed both ATL and PrC contribution to learning especially in the FMHO condition. We suggest that this ATL and PrC involvement might be relevant for different cognitive operations within the FM paradigm. To our understanding, learning by means of FM is not to be comprehended as a distinct learning *mechanism* but rather as an encoding *paradigm* comprising multiple mechanisms that contribute to rapid cortical integration of arbitrary associations. In particular, rapid cortical integration through FM includes the *discrimination* between pictures of complex objects, *binding* the visual features of an unknown item with an unfamiliar label to a feature conjunction that is represented as a coherent unit, and the *integration* of this unit into cortical memory networks. Considering the functional characteristics of the PrC and the behavioral findings of successful rapid learning through FM when high feature overlap item pairs were used (Sharon et al., 2011; Zaiser et al., 2019b), our approach was to increase the demands on the *discrimination* between the unknown and the known item. This should especially recruit the PrC, which might automatically trigger PrC-mediated *binding* mechanisms. We can imagine that successful discrimination and binding fosters cortical semantic integration, which in turn might be reflected by the ATL contribution to learning especially in the FMHO condition. Additional support of this idea of ATL-mediated semantic integration processes in the FMHO condition is provided by the way we manipulated feature overlap. The manipulation of visual feature overlap in our materials inevitably led to the simultaneous manipulation of semantic overlap. Although our intention originally was to manipulate visual overlap, we are aware that the current study cannot finally disentangle if it is visual or semantic overlap that drives the subsequent memory effects in the FMHO condition, especially as Martin, Douglas, Newsome, Man, and Barense (2018) recently reported that feature conjunctions are processed by PrC not only on a visual and a semantic level but also on an integrative level of visual and semantic features in combination. It is conceivable that an increase in both visual and semantic overlap could lead to greater PrC contribution to learning by means of FM, and that increased semantic overlap might have been additionally beneficial for the semantic integration process. The latter might potentially be mediated by ATL engagement, which would be in line with current models of the ATL as a semantic hub (e.g., Lambon Ralph et al., 2017).

It has been reported previously that fast and direct consolidation of new information is also possible if this information is congruent with a certain schema (e.g., Tse et al., 2007; van Kesteren et al., 2013; see van Kesteren, Ruiter, Fernández, & Henson, 2012, for a review). Schemas can be understood as higher-level structures of prior knowledge to which new information can be related. The embedding of new information into an existing schema can be facilitated if this information is congruent with the schema (see Gilboa & Marlatte, 2017, for a review). The benefit of schema congruency has been associated with mPFC involvement (van Kesteren et al., 2012; van Kesteren et al., 2013). In the present study, we also observed greater mPFC subsequent memory effects in the FMHO condition compared to the FMLO condition. Although the mPFC has been associated with many cognitive functions other than schema learning, the stronger mPFC memory contribution in the FMHO condition may reasonably be attributed to the stronger pre-activation of the relevant schema by the highly similar known item. For example, at encoding of the bird *satellote*, an accompanying flamingo likely has triggered the facilitating bird schema more strongly than a guinea pig. However, it is not yet clear if mPFC recruitment incrementally contributes to rapid cortical integration through FM or if it is rather a by-product that does not add to the contribution of other components such as the PrC and ATL, which might already be sufficient. More importantly, although schema-based learning might foster especially the integration of the picture of the unknown item, which is schema-congruent with the known item in the FMHO condition due to their strong semantic relation, it is unclear how schema-based learning alone could account for the binding of the unknown item and the arbitrarily matched (i.e., schema-incongruent) label.

In sum, we suggest that enhancing the demands on PrC involvement, operationalized by increasing feature overlap (as in the FMHO condition), supports learning by means of FM. ATL involvement may comparably foster learning in the FMHO condition, which we attribute to a stronger integration process. Furthermore, the potentially greater schema-congruency in the FMHO condition might have additionally contributed to the FM learning process, although current models of schema learning have difficulties to explain the binding of the unknown item to the label (see e.g., van Kesteren et al., 2012). However, the exact contribution of different cognitive operations and the underlying neurofunctional mechanisms driving rapid cortical integration in the highly complex FM paradigm yet needs to be further investigated.

The phenomenon of FM has so far been discussed predominantly with respect to the role of the hippocampus in memory for associations. It has been suggested earlier that learning by means of FM is hippocampus-independent (e.g., Sharon et al., 2011). The findings reported by Sharon et al. (2011) indeed suggest that hippocampal processing can be bypassed in learning of associations through FM since individuals with severe damage to the hippocampus were able to rapidly acquire novel associations. However, others reported contradictory findings. For example, no memory benefit from FM was observed for older adults with reduced hippocampal volume as a result of healthy aging (Greve et al., 2014) and hippocampal contribution to learning through FM in healthy young adults has been reported by Atir-Sharon et al. (2015) or at least could not finally be ruled out by Merhav et al. (2015). As already proposed previously (e.g., Atir-Sharon et al., 2015, and Merhav et al., 2014, 2015; Zaiser et al., 2019b), it might be over-simplified to claim that FM encoding is necessarily hippocampus-independent and hippocampal contribution to FM learning should be discussed in a more differentiated manner. First, recent research suggests that the hippocampus should not be considered a functionally homogeneous structure but might rather exhibit differences in functionality along its longitudinal axis. Whereas fine-grained recollection-like and navigational processes are allocated to more posterior parts of the hippocampus, the anterior hippocampus is associated with more coarse, gist-like representations, receiving schematic information from the ATL and object information from the PrC (e.g., Brunec et al., 2018; Brunec et al., 2019; Poppenk & Moscovitch, 2011; see Poppenk, Evensmoen, Moscovitch, & Nadel, 2013, for a review). This fits with the model of two cortical systems for memory-guided behavior by Ranganath and Ritchey (2012), suggesting that anterior parts of the hippocampus belong to the same anterior temporal system as the PrC and the ATL and are associated with semantic representations of objects rather than recollection-like retrieval and the tracking of episodic contexts. Notably, in the current findings, specifically anterior parts of the hippocampus contributed to learning by means of FM. This might possibly indicate that the anterior hippocampus as part of an anterior temporal system indeed plays a role in learning through FM, which might have been neglected so far. In previous discussions on the contribution of the hippocampus to learning within the FM paradigm, the definition of the hippocampus as a functionally homogeneous structure might not have been precise enough. Hence, we suggest that lesions of patients in studies on FM learning should especially be controlled for gradients along the longitudinal axis of the hippocampus.

The second important issue in the debate on hippocampal contribution to learning through FM is that the observed benefit for patients who cannot rely on hippocampal processing (see Sharon et al., 2011) does not allow for the reverse conclusion that FM is necessarily always independent of the hippocampus. In such patients who are unable to functionally rely on the hippocampus due to severe hippocampal damage, an alternative route triggered by FM encoding might make it possible to completely bypass hippocampal processing. However, this does not necessarily equally apply for other samples. For example, the conditions are less clear in healthy participants. The typically observed hippocampal degradation in healthy aging is also associated with a decline in learning of arbitrary associations compared to item memory (e.g., Naveh-Benjamin, 2000; Naveh-Benjamin, Hussain, Guez, & Bar-On, 2003). Yet, it is unclear to what extent the (only partly dysfunctional) hippocampal route is triggered by FM encoding in healthy older individuals. Greve et al. (2014) reported no learning benefit from FM in healthy elderly as measured by explicit recognition accuracy and even a positive relationship between the individual hippocampal grey-matter volume and recognition accuracy (in both the FM and EE condition), indicating that hippocampal processing might indeed play a role in learning through FM in healthy adults. However, analyses were not conducted separately for anterior and posterior parts of the hippocampus. Furthermore, it is conceivable that even if healthy elderly are impaired in hippocampal learning of associations, the route by which hippocampal processing could be bypassed was also not sufficiently triggered in Greve et al. (2014) as it was not controlled for feature overlap.

Apart from the finding that anterior and medial temporal lobe structures seem to contribute to learning especially within the FMHO condition, it is unclear which processes drove learning in the FMLO condition. One could assume that analogously to anterior hippocampal contribution in the FMHO condition, posterior parts of the hippocampus might have been involved more strongly in learning in the FMLO condition. Although this was not observed in the present experiment, the finding that the entorhinal cortex was engaged more strongly at retrieval in the FMLO condition might possibly be an indirect indicator of hippocampus-associated learning. Importantly, one should be aware that not finding the posterior hippocampus to be engaged in learning or retrieval in the FMLO condition does not allow for the conclusion that it has not been involved but might as well result from a lack of power.

## 5. Conclusion

Consistent with previous findings showing that feature overlap moderates rapid semantic integration after FM encoding on a behavioral level (Zaiser et al., 2019b), we conclude from the present results that differential PrC recruitment at encoding essentially influences rapid learning of novel associations within the FM paradigm. Beside the PrC, other anterior and medial temporal structures (i.e., the ATL and anterior hippocampus) were found to contribute to learning within the FM paradigm especially if the demands on object discrimination were high (operationalized by increasing feature overlap). Together with current knowledge of the functional characteristics of these brain structures, these findings provide further insights into the neurofunctional processes involved in FM. In future work, it would be interesting to investigate if triggering cognitive operations other than object discrimination can lead to a similar outcome.

## Acknowledgments

This work was funded by the German Research Foundation (grant numbers BA 5381/1-1 and ME 4484/1-1) in partial fulfilment of A.-K. Z’s doctoral dissertation. We would like to thank Luca Tarantini, Henry Schirok, and Laura Port for their help with data collection.

